# Interplay of Light, Melatonin, and Circadian Genes in Skin Pigmentation Regulation

**DOI:** 10.1101/2024.07.22.604624

**Authors:** Gabriel E. Bertolesi, Nilakshi Debnath, Neda Heshami, Ryan Bui, Hadi Zadeh-Haghighi, Christoph Simon, Sarah McFarlane

## Abstract

**Highlights:**

- Circadian pigmentation of tadpoles in vivo is mainly driven by melatonin
- Light and melatonin differentially regulate proliferation
- Melatonin mimics the expression of circadian core genes in the dark phase
- Deregulation of the circadian rhythm inhibits melanin synthesis

Circadian regulation of skin pigmentation is essential for thermoregulation, UV protection, and synchronization of skin cell renewal. This regulation involves both cell-autonomous photic responses and non-cell-autonomous hormonal control, particularly through melatonin produced in a light-sensitive manner. Photosensitive opsins, cryptochromes, and melatonin regulate circadian rhythms in skin pigment cells. We studied light/dark cycles and melatonin coordination in melanin synthesis and cell proliferation of *Xenopus laevis* melanophores. *In vivo*, tadpole pigmentation shows robust circadian regulation mainly hormone-driven, in that isolated melanophores respond strongly to melatonin but only slightly to light. Melanophore proliferation is faster in the dark and slower with melatonin compared to a 12/12 light/dark cycle. Expression of circadian core genes (clock, bmal1, per1, per2, per3, cry1, cry2, and cry4) in melatonin-treated cells during the light phase mimics dark phase expression. Individual Cry overexpression did not affect melanisation or cell proliferation, likely due to functional redundancy. Melanin synthesis was inhibited by circadian cycle deregulation through: a) pharmacological inhibition of Cry1 and Cry2 degradation with KL001, b) continuous light or dark conditions, and c) melatonin treatment. Our findings suggest that circadian cycle regulation, rather than proliferative capacity, alters melanisation of melanophores.

**Significance:** Circadian rhythms are a highly conserved phenomenon in nature. In vertebrates, the modification of skin pigmentation and epidermal cell renewal in response to the environmental light-dark cycle are crucial physiological adaptations that serve various purposes, including thermoregulation, reducing ultraviolet damage, and regulating skin stem cell proliferation. Our observations indicate that, *in vivo*, the circadian regulation of skin pigmentation is more influenced by cycling-melatonin levels than light/dark. The deregulation of the circadian cell cycle through various mechanisms all inhibited melanisation while cell proliferation was increased or reduced, suggesting that proliferation and melanisation are mechanistically dissociated responses.

Graphical Abstract

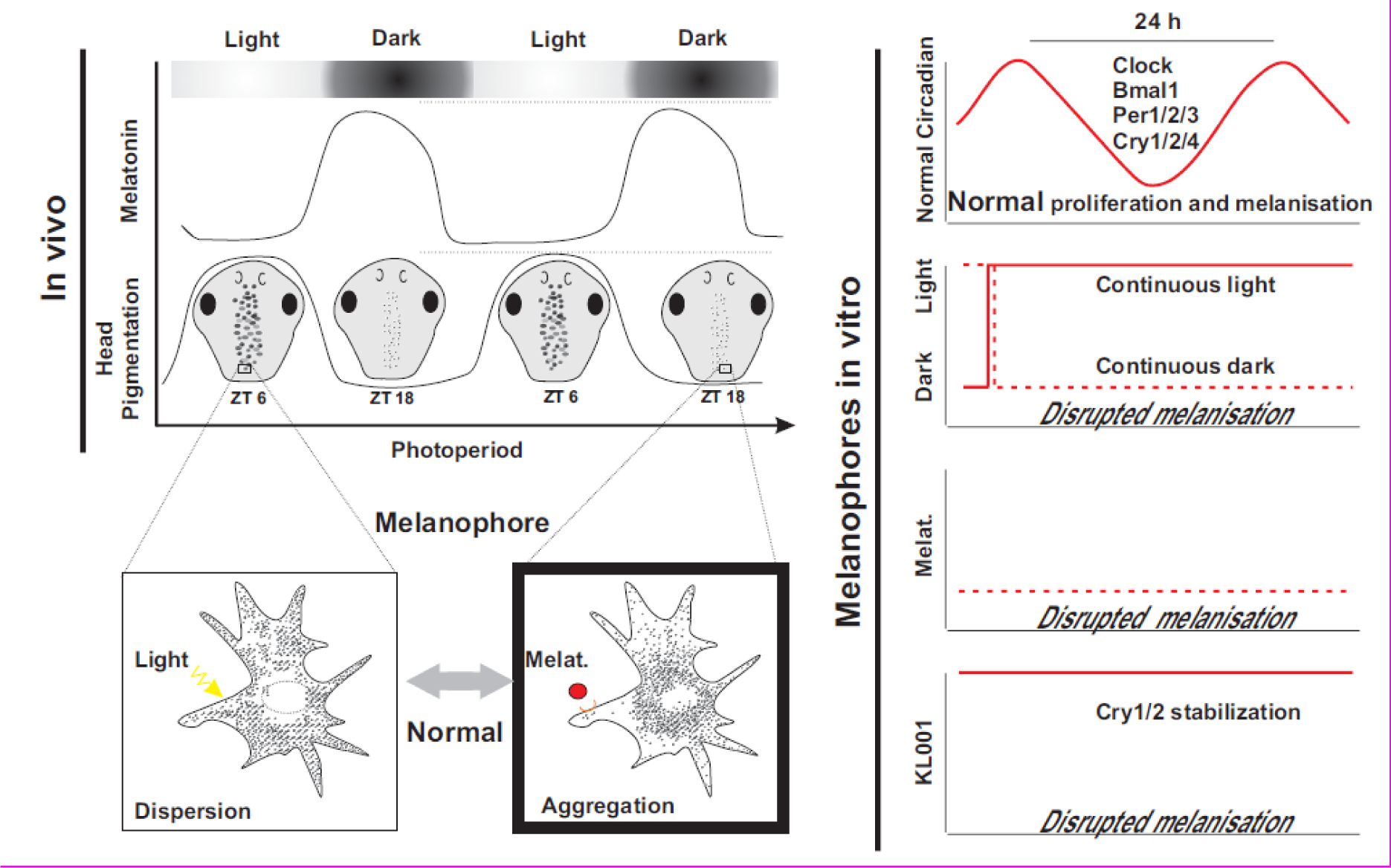

## 1. Introduction

Circadian rhythms are a highly conserved phenomenon in nature. In vertebrates, the modification of skin pigmentation and epidermal cell renewal in response to the environmental light-dark cycle are crucial physiological adaptations that serve various purposes, including thermoregulation [1], reducing ultraviolet (UV) damage [2], and regulation of skin stem cell proliferation [3]. In mammals, the cells that synthesize melanin pigment are called melanocytes and are evolutionarily related to the melanophores found in amphibians, reptiles, and fish. A combination of light-mediated cell-autonomous and neuroendocrine-hormone mechanisms trigger changes in skin physiology that include variation in pigment distribution/secretion and pigment cell proliferation. The cell autonomous physiological response is mediated by the direct photosensitivity of pigment cells through their expression of opsins [4, 5, 6]. Additionally, skin physiology is regulated cell non-autonomously by neuroendocrine hormones. Here, light works through opsins expressed by photosensing neurons of the eye and/or brain to control the circadian synthesis and release of melatonin, which affects both pigment distribution and cell proliferation [7, 8, 9]. How these two mechanisms are coordinated at the level of pigment cell behaviour is unclear.

Light is a prominent environmental signal that regulates the phase of the circadian clock and aligns the clock in individual cells with the ambient light. The skin can transmit light information to directly impact the circadian rhythm and proliferation of the epidermis. Light generally decreases proliferation: Light decreases epidermal stem cell numbers in S-phase of the cell cycle during the day compared to the night [10], reduces proliferation of cultured human melanocytes [11, 12], and inhibits proliferation of pigmented cells in fish [13]. Opsins such as Opn3, Opn4 (Melanopsin), and Opn5 function as light sensors in mammalian pigment cells to regulate both pigment synthesis/intracellular movement [4, 5, 6] and melanocyte proliferation [14, 15].

Light also controls skin pigmentation by regulating melatonin synthesis and secretion [8, 9, 16]. Melatonin inhibits both the synthesis of the pigment melanin and proliferation of melanocytes [12, 17]. Further, melatonin, secreted by the pineal complex when light levels drop below a minimum threshold, aggregates melanosomes in melanophores to lighten the skin of *Xenopus laevis* tadpoles [7,8]. In mammals, the melatonin-associated neuroendocrine mechanism plays the dominant role in the light-mediated control of circadian regulation of skin physiology. Indeed, melanocytes in fur-covered mammals reside deep within hair follicles so fail to experience the light signals faced by the integument of fish, amphibians, and reptiles. Instead, in mammals and birds, melanocytes transfer melanin granules to keratinocytes to pigment the hair and feathers [9, 18].

Cells in the different skin compartments of mice have intrinsic clocks that regulate their proliferation and differentiation, with external light sensed by intrinsically photosensitive retinal ganglion cells (ipRGCs; melanopsin-expressing cells) of the eye influencing these clocks via a neural circuit involving the central circadian pacemaker cells of the suprachiasmatic nucleus (SCN) [19, 20]. For instance, light uses this mechanism to stimulate the proliferation of murine hair follicle stem cells [20]. Whether the “lack” of light (dark) and melatonin, however, are interchangeable in the way they influence different skin pigment cell parameters, including proliferation, melanosome localization, and differentiation (melanisation) is unclear [17, 20]. Indeed, the influence of the direct and indirect (melatonin) light-mediated signals may vary among vertebrates, with organisms with uncovered integuments (fish, amphibians, and reptiles) relying on the former, and those with covered integument (birds and mammals) relying on the latter.

For both direct and indirect light-mediated mechanisms, light controls the circadian cycle by regulating the expression of four core clock genes that encode transcription factors: Period (PER), Cryptochrome (CRY), BMAL1, and CLOCK [21]. CLOCK and BMAL1 are transcriptional activators, while PER and CRY suppress gene expression [22]. Levels of the core genes are entrained or synchronized to the day-night cycle [22], and oscillate over the light/dark cycle in mammalian melanocytes [23, 24] and amphibian melanophores [25, 26]. PER1, BMAL1, and CLOCK regulate mammalian melanocyte pigmentation [27, 28], while melatonin and light coordinate core gene expression in *Xenopus* melanophores [25, 26]. Opsins play key roles in this regulation [4–6]. Whether other light-sensing molecules, such as the cryptochromes (Crys), which participate in circadian rhythm regulation and in some cases mediate blue light photosensitivity by non-covalently binding to a Flavin Adenine Dinucleotide (FAD) chromophore [29], participate is unknown.

Crys are divided in three groups: i) The type I or the *drosophila* type Cry (dCry); ii) the type II CRYs (CRY1 and CRY2) found in mammals; and iii) the type IV Cry (Cry4) present in several lineages of Chordata, including fish, amphibians and certain sauropsids [30, 31]. The mammalian type II CRYs lack photosensitivity and evolved as light-independent transcription factors that regulate core clock gene expression. In contrast, dCry photosensitivity entrains the *drosophila* circadian clock, and Cry4 acts in pigeons as both an UV/blue light photoreceptor and a light-dependent magnetosensor [29, 31, 32].

Larval *Xenopus laevis* provide an excellent model to study the direct and indirect influence of light on skin pigmentation, with *in vivo* and *in vitro* experimental approaches available to distinguish between their respective influence [7, 8, 33, 34]. In *Xenopus*, the melanophores are directly exposed to light because of their hairless integument. Light can also act via the pineal gland, however, to shut down melatonin secretion and aggregate pigment granules [33, 34]. Evolutionary analysis suggests that three Crys should be expressed in the amphibian *Xenopus laevis*: Cry1, Cry2 and Cry4 [30, 31]. Here we explore how light coordinates the direct and indirect regulation of pigment synthesis/movement and pigment cell proliferation, and whether light-sensitive Crys participate. We find that the light/dark deregulation of the circadian cycle, melatonin treatment, and pharmacological inhibition of the circadian degradation of Cry1 and Cry2 with KL001, all inhibit melanin synthesis. These data argue that the biosynthesis of melanin is linked to the circadian rhythm and is independent of the proliferative capacity of the cells.

## 2. Materials and Methods

### 2.1 *Xenopus laevis* tadpoles and assessment of pigmentation index

*Xenopus laevis* tadpoles were obtained by induced egg production from chorionic gonadotrophin (Intervet Canada Ltd.) injected females and *in vitro* fertilization according to the standard procedures (See protocol at Xenbase (http://www.xenbase.org; RRID:SCR_003280). Stage 43/44 embryos (staged according to Nieuwkoop and Faber [Xenbase (http://www.xenbase.org)] were used to determine circadian variation in skin pigmentation. The Animal Care and Use Committee, University of Calgary, approved procedures involving frogs and embryos (protocol# AC21-0148). Embryos were maintained at 16 °C until stage 24/26 (48 h), and then reared at room temperature (22–24 °C) until stage 43/44 in Marc’s modified Ringer’s (MMR) solution (100 mM NaCl, 2 mM KC1, 2 mm CaCl_2_, 1 mm MgCl_2_, 5 mM HEPES pH 7.4), under light cycles of 12 h ON (∼800 lux or 1.5 × 10–4W/cm^2^) /12 h OFF (∼ 2 lux). The skin pigmentation at different ZT times was analysed as described previously [35]. Briefly, pictures of the dorsal head of tadpoles were taken using a stereoscope (Stemi SV11; Carl Zeiss Canada, Ltd., Toronto, Canada) and a camera (Zeiss; Axiocam HRC), with identical conditions of light, exposure time and diaphragm aperture. Pictures were converted to binary white/black images using NIH ImageJ (U. S. National Institutes of Health, Bethesda, MD) public domain software and the density/number of positive pixels was measured. Significance (p < 0.05) between experimental groups was determined with GraphPad Prism 10.1 by using multiple ANOVA followed by Bonferroni’s post hoc test or t-test (n ≥ 10; N = 3).

### 2.2 *Xenopus* melanophore cell (MEX cell) culture

MEX cells are melanophores isolated initially from the ventral tail of *Xenopus laevis* tadpoles and established as a stable cell line [36]. MEX cells were obtained from Dr. Vladimir Rodionov at the University of Connecticut. Cells were cultured in 60% Leibovitz L15 medium, 35% water and supplemented with 5% heat inactivated fetal bovine serum (FCS) (named growing media here after) (Gibco, Canada, Lot #2635911RP) without antibiotics as described previously [33, 36]. Cells were grown at room temperature at the same light/dark cycles described above for tadpoles. Of note, medium without phenol red was used though all experiments to expose cell cultures to a visible spectrum of light radiation.

### 2.3 Determination of the population doubling time (PDT)

MEX cells were seeded at approximately 5×10^5^ cells/ml in a 35 mm^2^ grided dish in L15 growing medium. After 24 h, cells were kept for 10 days in constant darkness (<2 lux; DD), constant light (∼800 lux; LL), or in cycles of 12 h of light and 12 h of darkness (LD), either with control solution or with melatonin (10 nM) (Sigma) added every 48 h with medium changes at ZT=0. Bright field images of the same region of interest (ROI), as determined by the grid of the dish, were taken every day during the first hour of the light phase (ZT 0-1) for 10 days. Of note, the process of taking pictures required approximately 5 minutes/dish, therefore the DD-treated samples were exposed during that time to the light of the microscope. The number of cells in each ROI (n=8) was counted and expressed normalized to the number of cells at day one. Cell proliferation curves in each independent ROI approximated to a third order polynomial (cubic) regression. The normalized data of two independent experiments (N=2) were analyzed. The population doubling time (PDT) was calculated between day 1 and day 10 using PDT= T ln2/ln (X_1_/X_10_); T is the incubation time in hours (240 h; 10 days) and X_1_ and X_10_ are the number of cells in each area at the beginning and the end of the incubation time, respectively. Comparative analysis of the PDT was performed by One-way ANOVA, followed by Bonferroni’s multiple comparation using the GraphPad Prism (10.1.2) software. p<0.05 was considered statistically significant.

### 2.4 Melanin synthesis assay

MEX cells treated with phenylthiourea (PTU; 1 mM), an inhibitor of melanin synthesis [37], were seeded in a 35 mm dish in L15 growing media. Cell growth and melanin synthesis was monitored by taking, for approximately 15 days, daily pictures of the dishes during the first hour of the photophase (ZT=0 to ZT=1). A densitometric analysis of the pigmented dish was determined using the ImageJ software, as described previously for tadpoles [35]. Biochemical determination of melanin synthesis was performed as described previously [38]. Briefly, melanophores were harvested, and the number of cells quantified in a Neubauer counting chamber. Resuspended cells (3 x 10^5^ cells / ml) were lysed in phosphate buffered saline (PBS) containing 1% Triton X-100. After centrifugation, the melanin-containing pellet was resuspended in 1N NaOH and incubated at 80 °C until fully dissolved to measure absorbance at 405 nm in a spectrophotometer. Absorbance data are expressed as % of control (n=4; N=2).

### 2.5 RNA isolation, cDNA synthesis, cloning of cryptochromes and generation of stable cell lines

Total RNA was extracted from MEX cells (sub confluent monolayer in a 35 mm dish) using the TRIzol reagent (Invitrogen) method and following the protocol provided by the manufacturer. Equal volume of 100% ethanol was added to the upper aqueous layer from Trizol-centrifuged samples and the RNA was purified using purification kit columns (Thermo Scientific GeneJET RNA Purification Kit). RNA was then treated with DNAse-I and eluted using a nuclease column (BioRad) and RNAse free water. Total RNA extracted from MEX cells was converted to cDNA using the SuperScript^TM^ IV First-Strand Synthesis System (Invitrogen) using oligo(dT) primers to prime the samples according to the manufacturer’s protocol. Specific primers were designed to amplify both the L (long) or S (short) homologous gene from the tetraploid *X. laevis* specie [39] (Supplementary table 1). All RT-PCR amplifications were carried out in a total volume of 20 μL with 1 μL of cDNA, 2 μL of primers, 7 μL of water and 10 μL of 2X PCR master mix (Thermo Scientific, IL). PCR products obtained from cDNA were cloned into TOPO-pCRII (Invitrogen) vectors and sequenced at the DNA Services Facility, University of Calgary. The NCBI gene access number of the cloned construct correspond to NM_001087660 (nucleotide 160 and 2230), NM_001090467.1(nucleotide 97 to 1900) and XM_018245284 (nucleotide 33 to 1779) for *cry1.L*, *cry2.S* and *cry4.L,* respectively. Cry1/2 and 4 were individually subcloned into a pCDNA3.1 vector to obtain stable cell lines. MEX cells were transfected with Lipofectamine 2000 (Thermo-Fisher Scientific) according to the manufacturer’s protocol. The plasmid pcDNA3.1-Vector (Vect) or pcDNA3.1 expressing the full-length *cry* genes, pcDNA3.1-*cry1*, pcDNA3.1-*cry2* or pcDNA3.1-*cry4,* were transfected to generate the respective stable cell lines. Cell selection was performed with 750 µg/ml of G418 (Roche) for approximately 4 weeks until individual clones were isolated and tested for *cry* overexpression by RT-PCR.

### 2.6 Immunohistochemistry

Immunohistochemistry was performed on 12 µm sections of stage 44/45 tadpoles or in MEX cells grown on Nunc® Lab-Tek® II Chamber Slide™ system, which were fixed with 4% paraformaldehyde in the middle of the light phase. The samples were then washed with PBS once and then 5 times with PBT (0.2% Tween-20 in PBS). The samples were blocked using a PBS/Tween 0.2% solution containing 5% goat serum for 30 minutes. Anti-clock primary antibody (Life Span BioSciences; rabbit polyclonal against human clock; 1/200 dilution; LS-C109824) or anti-cryptochrome 1 (MyBioSource; MBS3214492) in blocking solution were placed on the sample for 90 minutes. Anti-rabbit Alexa Flour 555 (Invitrogen; 1:1000 dilution) was used for detection. The nuclei were stained with Hoechst 33243 (1 µg/ml dilution in PBT). All incubations were conducted at room temperature. Aquapolymount was used to mount slides.

### 2.7 Statistical analysis and figure preparation

Statistical analyses are described in the individual figure legends and in the method sections for each separate experimental assay. Significant statistical differences between treatments were analyzed with the GraphPad Prism 10.2.3. software and figures generated with GraphPad Prism and CorelDraw 10.1 software.

## 3 Results

### 3.1. *Xenopus* tadpole melanophores *in vivo* change their pigmentation with the dark-light circadian cycle; *In vitro,* however, a few cells respond to light but strongly respond to melatonin

The neuroendocrine system mediates a rapid (45 minutes), robust and quantifiable change in skin pigmentation when tadpoles are placed in the dark [7, 8, 40]. Melatonin released from the pineal gland during the night lightens the skin of stage 43/44 *Xenopus* tadpoles and adult frogs [8, 41], a process that is shut down by light acting via the pineal complex and not the eye [8]. To determine whether the light-dark cycle additionally regulates pigment localization via direct light actions on melanophores we compared the skin pigmentation response of *Xenopus* tadpoles over the full circadian cycle to that exhibited by melanophores in culture. First, we determined circadian pigmentation changes in *Xenopus* tadpoles *in vivo*. We maintained animals under a 12-hour light ON/12-hour light OFF cycle from the first day of oocyte fertilization until stage 45/46 (approximately 5-6 days at 20°C), and then measured the pigmentation index of the dorsal head as described previously [35]. We observed maximum pigment dispersion at zeitgeber (ZT) 6-8, and skin lightening, through pigment aggregation, in the dark phase from ZT12 to ZT24 (Fig. 1 A and 1 B).

**Figure 1:**
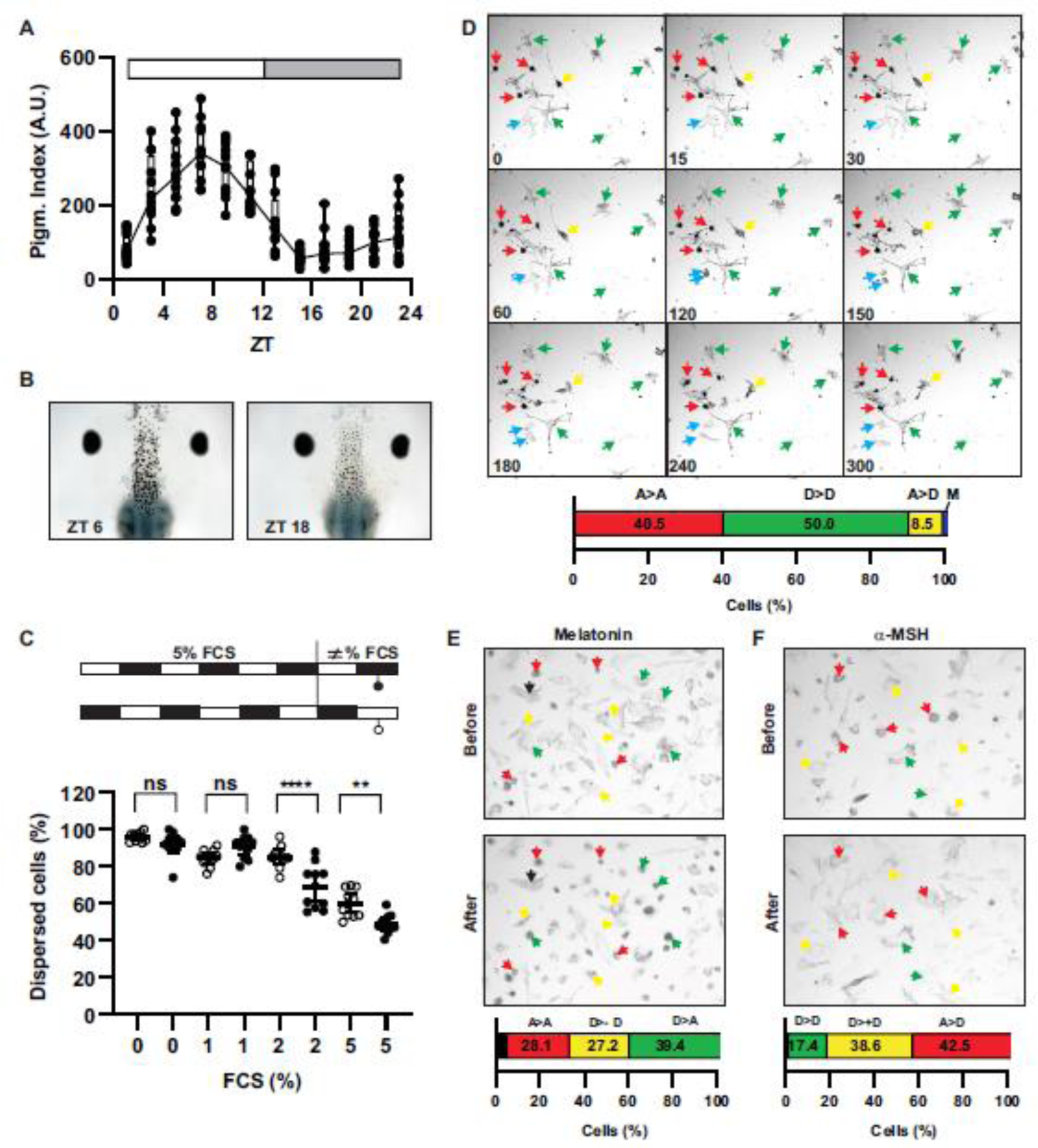
Circadian regulation of skin pigmentation *in vivo* and the effect of light and melatonin on pigment distribution *in vitro*. **A)** Pigmentation index of tadpoles at stage 45/46 as measured from the dorsal head: Tadpoles were maintained in a 12-hour light ON: 12-hour light OFF (L:D) cycle. Dots represent the pigmentation index for each tadpole, and rectangles indicate the median with 25-75 percentiles. **B)** Representative picture of the dorsal head of a tadpole at indicated Zeitgeber Times (ZT). **C)** Melanosome distribution state (Aggregated/Dispersed) of melanophore MEX cells *in vitro.* MEX cells were cultured in Light: Dark conditions as illustrated in the diagram. Various percentages of FCS were added in the last 24 hours. t-Test; **p<0.01; ****p<0.001. ns; nonsignificant. **D)** Kinetics of the response of MEX Cells exposed to light. The response of MEX cells to light exposure is depicted, showing cells that remain aggregated (A>A; red arrows), remain dispersed (D>D; green arrows), disperse their melanosomes (A>D; yellow arrow), or undergo cell division (M; mitosis; blue arrow). **E)** MEX cell melanosome distribution before and after 1-hour treatment with melatonin (10 nM) or αMSH (100 nM). The percentage of cells exhibiting different melanosome distributions, as indicated in D, plus unresponsive cells (dispersed or aggregated in melatonin or αMSH treatment, respectively; labeled black) is shown.

Next, we investigated whether cultured melanophores also respond to light. Previous studies indicate that light promotes melanosome dispersion of cultured melanophores [33, 34] and melanophores of isolated tails [42], however, the degree of the response, and a comparative analysis with the response induced by hormones have not been performed. We used MEX cells, a stable melanophore cell line obtained from *Xenopus* tadpoles [36]. Because melatonin and α-MSH are present in the blood [16, 43] and the fetal calf serum (FCS) used to supplement the cells, we tested different concentrations of FCS for their ability to aggregate/disperse melanosomes. MEX cells were cultured during five days with the same light intensity and cycles (12-hour light ON/12-hour light OFF) as previously performed with tadpoles, in L15 medium without phenol red supplemented with 5% FCS, and then switched to media with different FCS concentrations for 24 hours (Fig. 1 C, insert). The number of cells with aggregated/dispersed pigment in the middle of the light and the dark phase were counted (Fig. 1 C). The proportion of aggregated cells increased with a higher FCS concentration, reaching approximately 50% in semi-confluent cultures with 5% FCS (Fig. 1 C). Because we found the proportion of cells with aggregated vs. dispersed pigment varied between FCS batches, the results described here were obtained using the same FCS batch.

Interestingly, a small but significant increase was seen in the number of cells with dispersed pigment in the light phase vs. the dark phase at both 2.5% and 5% FCS (Fig. 1 C). These data suggest that light induces pigment dispersion in only a small proportion of cells. To understand how pigmentation changed in response to light on an individual cell basis, we performed time-lapse analysis of MEX cells for 5 hours after the cells were moved at the end of dark phase of the cycle (ZT 24) to continuous light exposure from the microscope, a stimulus that *in vivo* would cause dispersion. We identified four types of aggregation/dispersion responses: i) *aggregated cells insensitive to light*, which did not change pigment localization and remained aggregated (Fig. 1 D; red arrows; 40.5% (85/210) of cells; n=4; N=4); ii) *dispersed cells insensitive to light* that retained their dispersed characteristics (Fig. 1 D; green arrows; 50% (105/210) of cells; n=4; N=4); iii) *light-sensitive cells*, which switched from an aggregated to a dispersed state (Fig. 1 D; yellow arrow; 8.5% (18/210) of cells; n=4; N=4); and iv) a small number of *mitotic cells* (Fig. 1 D; blue arrows; 0.95% (2/210) cells n=4; N=4) that showed aggregated pigment during cell division.

We next compared how the light response compares to the pigment aggregation and dispersion triggered by melatonin and alpha melanocyte stimulating hormone (α-MSH), respectively. Interestingly, many fewer MEX cells responded to light (approximately 10%) than responded to melatonin or α-MSH. Most cells (66.6%) showed full or partial responsiveness to melatonin (10 nM) [(Fig. 1 E; green arrows; Dispersed to aggregated (D > A); 39.3% (130/331); n=4; N=4)] or partially aggregated (D > less D; D> -D) after 1 hour of treatment (Fig 1 E; yellow arrows; 27.2% (90/331); n=4; N=4). Further, approximately 28.1% (93/331) of the cells (Fig 1 E; green arrow) remained aggregated, and 5% (18/331; Fig 1 E; marked black) remained dispersed. MEX cells were also highly responsive to α-MSH (10 nM): 80% of cells responded to α-MSH, transitioning to a dispersed state [(Fig 1 F; red arrows; (A>D), 42.5% (100/249); n=4; N=4)] or increasing the level of dispersion [(Fig 1 F; yellow arrows; (D>+D) 38.6% (100/249); n=4; N=4)]; “No response” cells included cells that remained in their initially either dispersed (17.3%, 43/249) (Fig 1 F; green arrow; D>D) or aggregated (2%, 4/249) (Fig 1 F; marked black) state over the recording period.

Together, these results demonstrate that while tadpole melanophores exhibit a robust circadian response *in vivo*, where melanosomes disperse during the light phase and aggregate during the dark phase, the sensitivity of melanophores in culture to light is minimal. Yet, these cells show strong responses to α-MSH and melatonin. These results suggest that similarly *in vivo* there is little or no pigmentation response of melanophores directly to changes in light illumination.

### 3.2 Light and melatonin both alter melanophore proliferation and circadian core gene expression

Dark and melatonin both induce melanosome aggregation, in contrast to the dispersion seen with light. Additionally, melatonin and its metabolites inhibit mammalian melanocyte proliferation [12, 17, 44]. Thus, we asked whether melatonin and/or dark similarly impact *Xenopus* melanophore proliferation in culture. First, we examined the impact of light/dark on the proliferation of MEX cells subjected to different lighting conditions: constant darkness (DD), constant light (LL), and 12 hours light ON and 12 hours light OFF cycle (LD). Cultures maintained in the dark exhibited the fastest growth (Fig. 2 A), with a significantly reduced population doubling time as compared to cells kept in either continuous light or on a light/dark cycle (Fig. 2 B). In contrast, when we introduced melatonin (10 nM) at the beginning of the light phase (ZT0), to simulate continuous darkness, we observed less cell growth (Fig. 2 C), as seen as an increased population doubling time (Fig. 2 D). These data suggest that unlike what is seen with melanosome movements, cell division is differentially regulated by darkness and melatonin.

**Figure 2:**
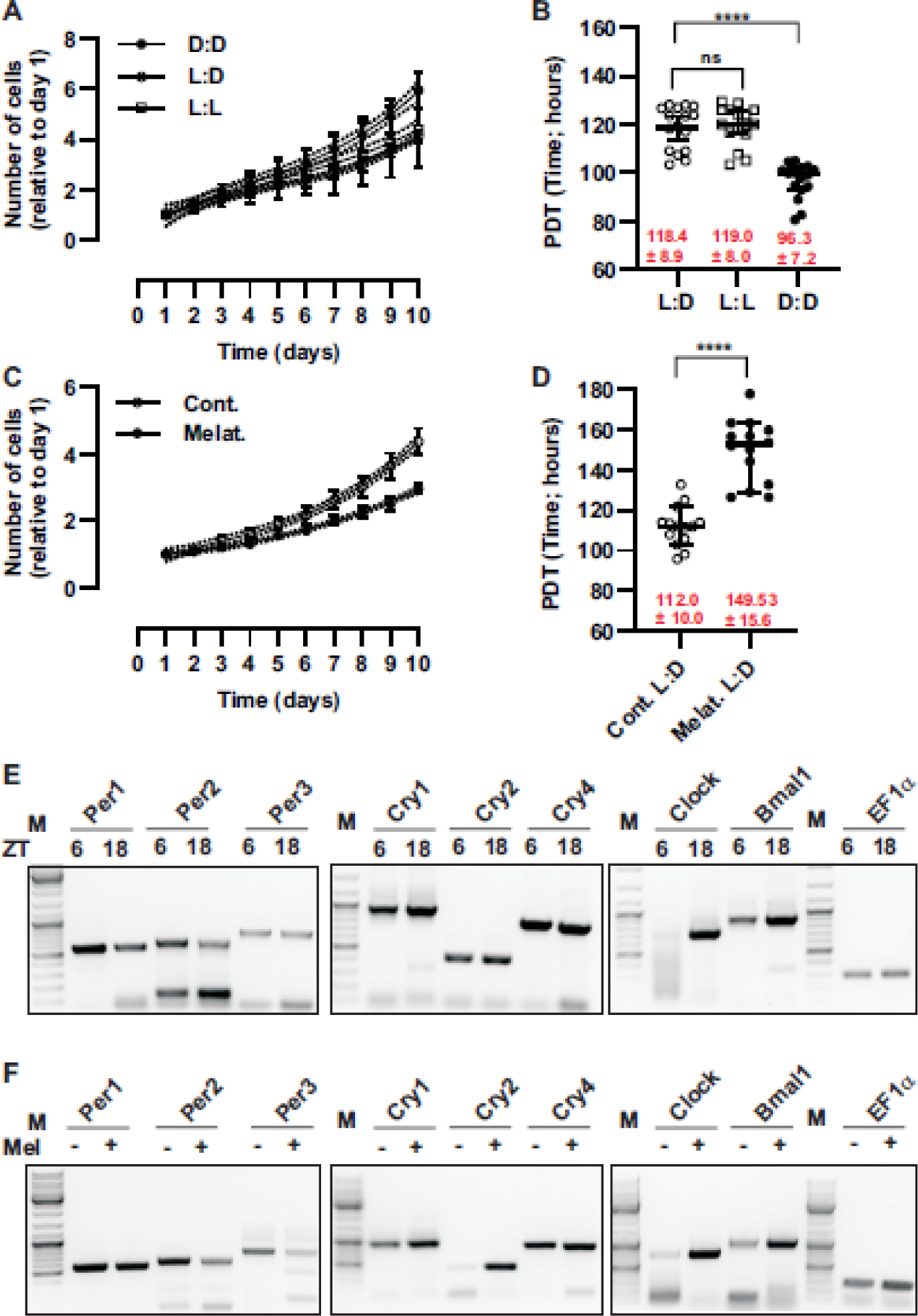
Proliferation of MEX cells and expression of circadian core genes are affected by light/dark and melatonin. **A, C)** Cell proliferation curve under different light conditions (A) or in the presence of melatonin (10 nM) (C): Cells were cultured in gridded dishes, and the number of cells in a specific region of interest (ROI) were counted daily over a 10-day period. Each data point represents the mean ± 95% confidence interval (CI) from 14 ROIs (n=14) in two independent experiments (N=2). Data are normalized to the cell count on day one. The proliferation curve is fitted to a third-order polynomial (cubic) regression, with the 95% CI indicated by dotted lines. **B, D)** Population doubling time (DPT) in hours of cells grown in different light conditions (B) or treated with melatonin (10 nM) (D); DPT was determined between day 1 and day 10. Data are presented as mean ± 95% CI; n=14; N=2. ****; p<0.0001 multiple ANOVA followed by Bonferroni (B) or t-Tests (D). **E, F)** RT-PCR analysis of circadian core genes in MEX cells. Expression of the indicated circadian core genes was measured during the middle of the light phase (ZT6) and dark phase (ZT18) (E), or after treatment with melatonin (10 nM) administered at the beginning of the light phase (ZT0) and measured at ZT6 (F).

We also compared the effects of dark and melatonin on melanophore expression of circadian clock genes. *Xenopus* melanophores express several core circadian genes, including *clock*, *bmal1*, *per1* and *per2* [25, 26]. We asked if MEX cells express these core genes, as well as additional core genes, *per3* and the three *cry* genes (*cry1*, *cry2*, and *cry4*), whose expression in melanophores is unknown. Of note, *X. laevis* is an allotetraploid species originating from diploid progenitors approximately 17–18 million years ago [39]. The duplicated genome of *X. laevis* is referenced to *X. tropicalis* but denotes an “L” or “S” for the longer and shorter homologous chromosomes, respectively. To identify which circadian core genes and type of homologue were expressed in MEX cells, we designed primers capable of amplifying genes on both the L and S chromosomes. Subsequent cloning and sequencing of the PCR amplicons revealed that *Xenopus* MEX melanophores predominantly express the “L” homologs, such as *per1.L*, *per2.L*, *bmal.L*, *cry1.L*, and *cry4.L*. The “S” expressed variants were detected for *per3.S* and *cry2.S* (Fig 2 E, 2 F and 4 A). We also detected Clock and Cry1 protein in *Xenopus* skin melanophores and MEX cells *in vitro* by immunohistochemistry (Suppl. Fig 1) using antibodies generated against the amino terminus of human CLOCK and CRY1, which possess high homology with the *Xenopus* proteins (Suppl. Fig 2). We next examined the regulation of these clock genes by light, comparing mRNA levels by RT-PCR at the middle of the light (ZT6) and the dark (ZT18) phase. As expected, *bmal1* and *clock* mRNA levels were higher in the dark than in the light phase, while *per1* and *per2* exhibited higher expression during the photophase than the scotophase (Fig. 2 E) [25, 26]. Interestingly, the transcriptional repressors *per* and *cry* were differentially regulated, with *per* and *cry1*/2 mRNA levels highest during the day and night, respectively (Fig. 2 E). Note that *cry4* showed minimal variation between light and dark phases

We then compared these light/dark levels of circadian core genes with those induced by melatonin. Cultures maintained for 5 days in light cycles of 12 h ON/12 h OFF were treated on the last day with melatonin at the beginning of the light phase (ZT0) and the expression of circadian core genes was analyzed 6 hours later (ZT6; middle of the light phase). Melatonin produced the same pattern of circadian core gene expression as observed at ZT18 (Fig. 2 F), suggesting that melatonin mimics night conditions.

Our results suggest that darkness and melatonin regulate similarly the expression of circadian core genes, with *per1* and *per2* mRNA levels highest during the day, and *cry1* and *cry2* mRNA levels peaking during the dark phase and in response to melatonin. Interestingly, however, these two signals have opposite effects on melanophore proliferation, decreasing and increasing the population double time, respectively.

### 3.3 Dark, light and melatonin treatment reduce melanisation of melanophores

We next studied how the light/dark cycle and melatonin impact melanisation (melanin synthesis). Light acts on opsins to activate pathways in cultured mammalian melanocytes associated with melanin synthesis. OPN3 [4], melanopsin (OPN4) [45], and OPN5 [5] are all implicated in melanogenesis. We showed previously that MEX cells express *opn5* and two melanopsin genes (*opn4* and *opn4b*) [46]. Of note, *opn4b* disappeared in the mammalian lineage during evolution [47]. To examine the impact of light/dark and melatonin on melanogenesis, we de-pigmented MEX cells by treatment with phenylthiourea (PTU), an inhibitor of melanin synthesis [37]. Cells were then sub-cultured into 35 mm dishes and PTU removed to initiate melanin synthesis. The cells were subjected to either continuous light (LL), continuous dark (DD), or light-dark cycles (LD). Melanogenesis was assessed daily by analyzing the degree of pigmentation in the dish [48]. Over a 15-day period, melanogenesis appeared to increase more for cells under a LD cycle or maintained in continuous dark (DD) than those always exposed to light (LL) (Fig. 3 A and 3 B). Because cells proliferate more under dark conditions, our melanisation measures were likely overestimated. To more accurately determine the melanisation level, we resuspended the cells at the end of the 15-day experiment and measured melanin content. A suspension culture from DD conditions (∼3 x 10^5^ cell/ml) showed a significant (15%) decrease in their pigmentation when compared to the LD-treated cells (Fig 3 C). Cells kept under light conditions (LL) cells were 45% lighter than cells under LD cycles (Fig 3 C).

**Figure 3:**
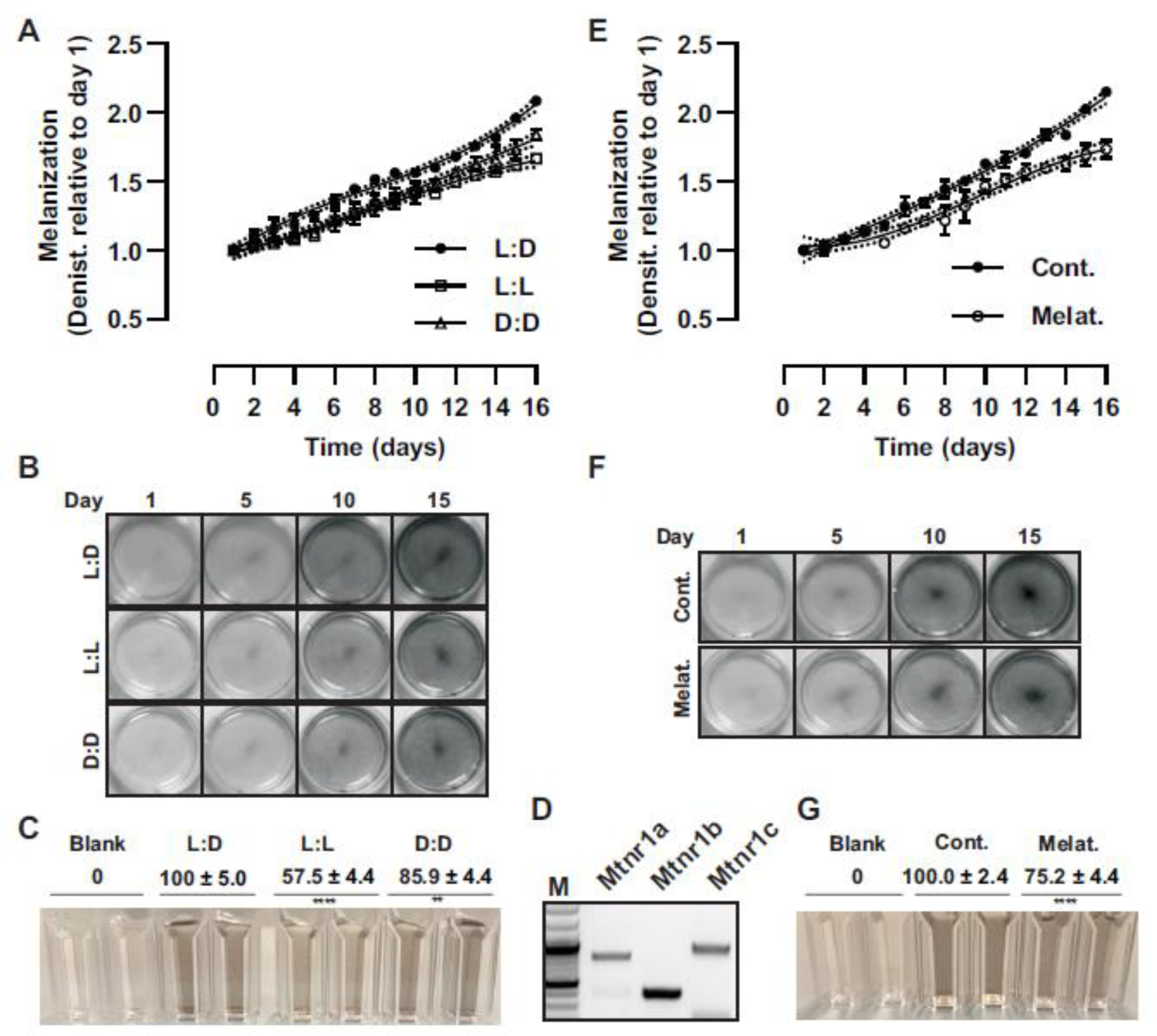
Dark, light and melatonin decrease melanisation. **A, E)** MEX cells depleted of melanin by PTU treatment were seeded in 35 mm dishes and maintained under light:dark cycles (L:D) 12 h light ON 12 h OFF, continuous dark (D:D), or continuous light (L:L) (A) or treated with melatonin (10 nM) (D). The effect on melanisation was determined by densitometric analysis of pictures obtained every day from the cell monolayers. Data from a representative experiment are presented as mean ± SD; n=4; N=2. **B, F)** Representative picture of dishes taken at the indicated days when treated under different light exposures (B) or with melatonin (F). Melatonin was added every two days at the beginning of the light phase (ZT=0). **C, G)** Cells in suspension (∼3 x 10^5^ cells/ml) after 15 days of the indicated treatments. Melanin content is expressed as relative to control. Data are the mean ± 95% CI; n=8; N=2. **, p<0.01; ****; p<0.0001 multiple ANOVA followed by Bonferroni or t-Test **D**) RT-PCR analysis of melatonin receptors expressed in MEX cells.

We next asked whether melatonin also impacted melanogenesis of MEX cells. Melatonin works on human melanocytes through melatonin receptors MTNR1A and MTNR1B (also known as MT1 and MT2) to inhibit both melanogenesis and proliferation [17]. Amphibians have three melatonin receptors, Mtnr1a, Mtnr1b, and Mtnr1c, with the *mtnr1c* gene disappearing during mammalian evolution [50]. We showed previously that *mtnr1a* and *mtnr1c* are highly expressed in melanophores of the tail of *X. laevis* [8]. Using RT-PCR, we detected the expression of all three melatonin receptors in MEX cells (Fig 3 D). We then analyzed the melanogenesis response in cells treated with melatonin. Similar to what was seen with dark conditions, the melatonin-treated cells exhibited reduced pigmentation as compared to controls (Fig 3 E and 3F). The inhibition induced by melatonin occurred early in the treatment (day 1 to 6) and persisted (Fig 3 E and F). In support, resuspended 15-day old cultures showed a 25% decrease in melanin content (Fig 3 G).

Together, our results suggest that darkness and melatonin have similar effects on melanisation and the regulation of circadian core genes, particularly in inducing the expression of *cry1* and *cry2,* yet differ in their effects on pigment cell proliferation.

### 3.4 Cry is not sufficient to modify cell proliferation, melatonin sensitivity, and melanisation

We find that continuous dark and melatonin treatment inhibit melanisation alongside an upregulation in *cry1* and *cry2* expression. Interestingly, Gao and coworkers [49] showed recently that KL001, a compound that blocks ubiquitination and photo-biomodulation of Cry1 and Cry2 [50, 51], hinders melanogenesis of mammalian melanocytes *in vitro* and *in vivo* [49]. Thus, we hypothesized that melatonin and the dark upregulate Cry to inhibit melanophore melanogenesis. If true, we predicted Cry overexpression would mimic melatonin’s effects on melanogenesis. To test this idea, and to identify whether a specific Cry was important, we first identified the full-length *cry* genes, *cry1.L*, *cry2.S*, and *cry4.L* from MEX cells (Fig 4 A). These genes were subcloned into the expression vector pcDNA3.1. Stable cell lines that overexpressed each construct or the empty pcDNA3.1 vector (Vector) were selected with G418 over a period of 2 months. Three distinct stable overexpression cell lines (A, B, C) were chosen for the vector and individual *cry* genes for subsequent analysis. Each stable cell line expressed mRNA for a specific *cry* (Fig 4 B). Notably the *cry2*-overexpressing cells showed a slight downregulation of *cry1* and c*ry4* (Fig 4 B).

**Figure 4:**
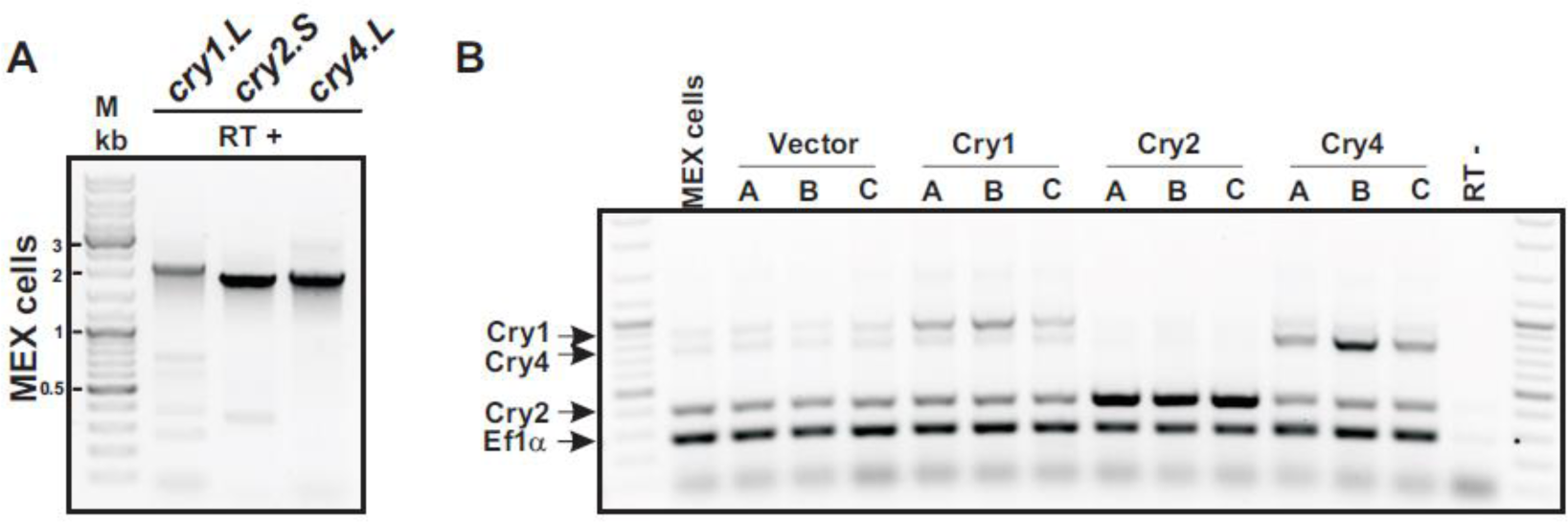
Generation of *cry* overexpression MEX cell lines. **A)** Full length *cry1.L*, *cry2.S* and *cry4.L* genes amplified by RT-PCR from MEX cells. **B)** RT-PCR analysis of *cry* genes in MEX cells and three stable cell lines (A, B or C) that overexpress either the pcDNA3.1 empty vector (Vector) or the full-length *cry* (*cry1*; *cry2* or *cry4*). Of note, specific internal primers were used to amplified simultaneously all *cry* genes and the housekeeping gene *ef1α*.

It was important to verify that the stable cell lines retained their responsiveness to melatonin and associated downstream signaling pathways. Cell lines were treated with melatonin for 1 hour, mirroring our previous experiments with the “maternal” MEX cell line (Fig 1 E), and pigment localization was assessed. By analysing the live effect of individual cells to melatonin we determined that all the cells responded to melatonin by pigment aggregation (Fig. 5 A). Indeed, after 1 hour of melatonin the combined percentage of partially responsive cells (initially dispersed to a less dispersed final state; D > -D; Fig. 5 B; yellow segment of the bar) and cells with completely aggregated pigment (dispersed to aggregated; D > A; green segment of the bar) ranged from 60% to 70% across all cultures (Fig. 5 B).

**Figure 5.**
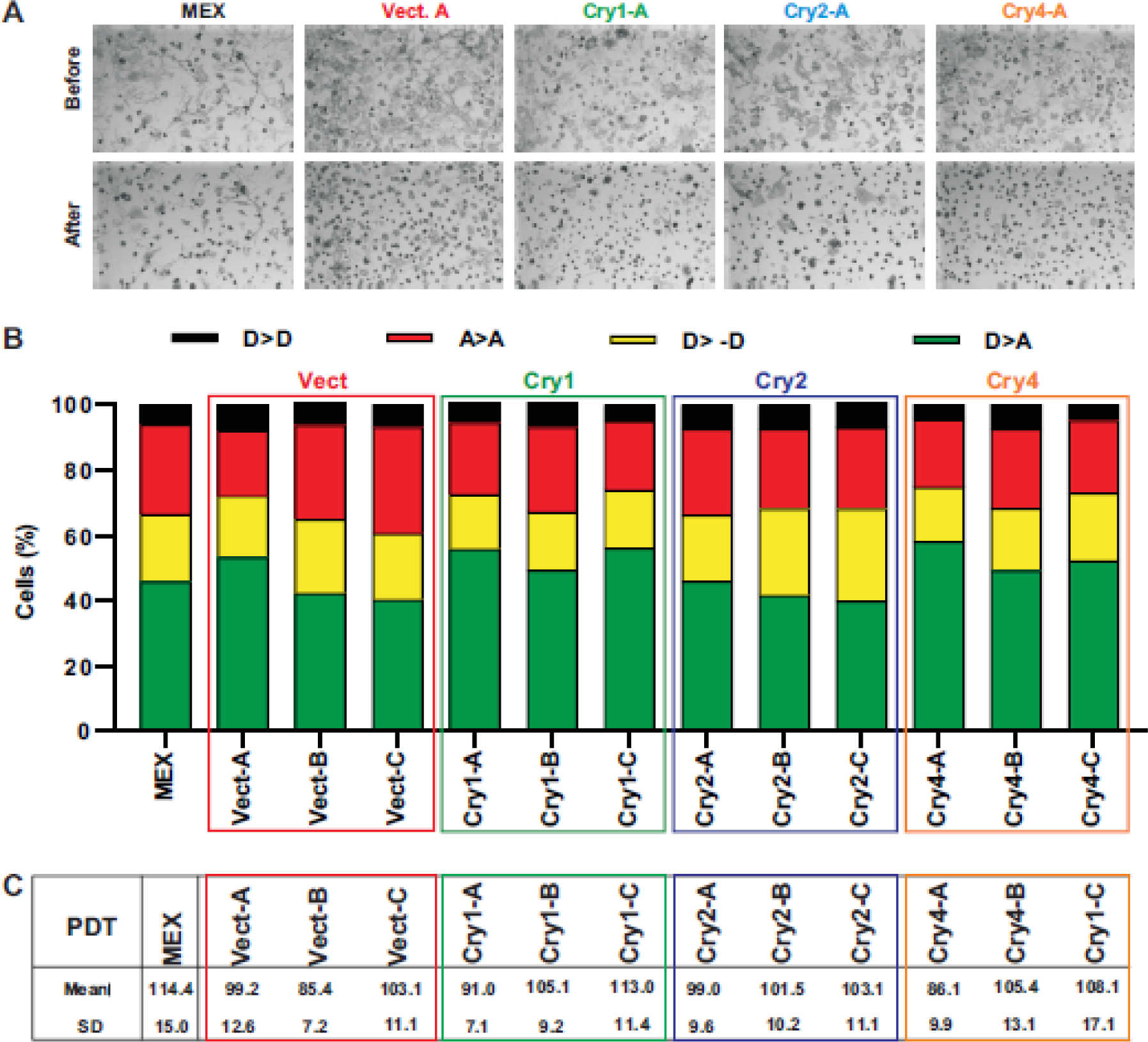
Cry overexpression does not affect melatonin sensitivity and cell proliferation. **A)** Melanosome distribution before and after 1-hour melatonin (10 nM) treatment of MEX cells and three stable cell lines (A, B or C) that overexpress either the pcDNA3.1 empty vector (Vector) or the full-length *cry* (*cry1*; *cry2* or *cry4*). **B)** The percentage of cells exhibiting different melanosome distributions is indicated, with cells that remain aggregated (A>A; red), remain dispersed (D>D; black), partially aggregate their melanosomes (D>-D; yellow), or completely aggregate melanosomes (D>A; green). **C)** Population doubling time (PDT) in hours of cells grown under Light: Dark conditions (12 h: 12 h). PDT was determined between day 1 and day 10 as indicated in Figure 2. Data are presented as mean ± 95% CI; n=12; N=2.

We next asked if Cry overexpression mimicked melatonin treatment, namely decreased population doubling time and inhibited melanogenesis. We measured the cell growth and the population doubling times under light: dark cycles. No significant differences were observed among the three sets of Cry-overexpression clones to those expressing the vector alone. Note that there were minor variations in the population doubling times between cell lines (Fig. 5 C), likely linked to clonal expansion within each cell line. We noticed that all selected clones, including vector controls, showed a slightly shorter population doubling time than the parental MEX cells, presumably because of the presence of G418 in the culture media.

Cry overexpression also did not appear to impact the synthesis of melanin. Similar increases in melanisation with time in culture were observed for the maternal MEX cells, the clones containing the empty vector, and those overexpressing Cry1, Cry2, or Cry4 (Fig 6 A - E). Together, these observations suggest that Cry is not sufficient to alter the melanisation or proliferation of melanophores.

**Figure 6.**
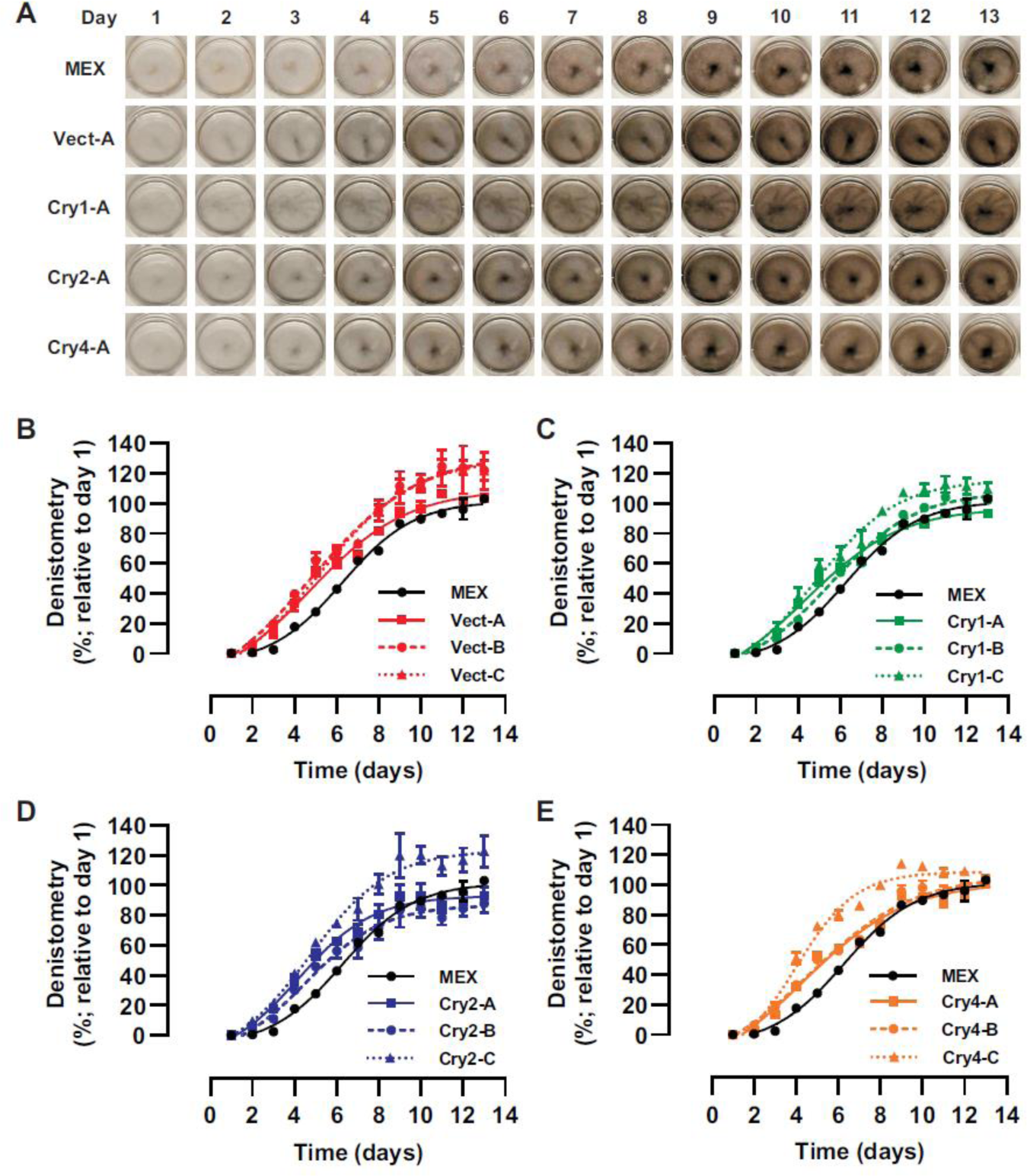
Cry overexpression does not affect melanin synthesis. **A)** MEX cells and stable clones overexpressing Crys were depleted of melanin by PTU treatment and seeded in 35 mm dishes as indicated in Figure 3. Cells were maintained under light: dark cycles (L:D) (12 h light ON 12 h OFF). Only clone “A” is shown. **B-E)** Pigmentation level determined by densitometric analysis of pictures obtained every day from cell cultures comparing the parental MEX cells (black line) against control stable line (Vector) (B) or overexpressing the Cry1 (C), Cry2 (D) or Cry4 (E). Data are the mean ±SD from one representative experiment.; n=2; N=3.

### 3.5 Pharmacological biomodulation of cry inhibits melanisation

To test the necessity of Crys in dark-induced increases in pigmentation we used KL001, a non-selective molecule that stabilizes both Cry 1 and Cry2 to lengthen the circadian period [50]. KL001 inhibits melanization of B16 melanoma cells [49]. To test differential requirements of Cry1 and Cry2 we also used KL101, a compound that selectively stabilizes Cry1 with no effect on Cry2 [52]. Of note, whether either of these small molecules stabilizes Cry4 is unknown.

Treatment with KL001 (10 µM) dramatically reduced melanisation of MEX cells, while KL101 (10 µM) showed a partial inhibition (Fig. 7 A-C). These results indicate that Crys participate in melanisation. The involvement of both Cry1 and Cry2 is suggested by the observation that while photo-stabilization of both molecules results in a strong decrease of melanin synthesis, only a partial response is present when only Cry1 is stabilized.

**Figure 7.**
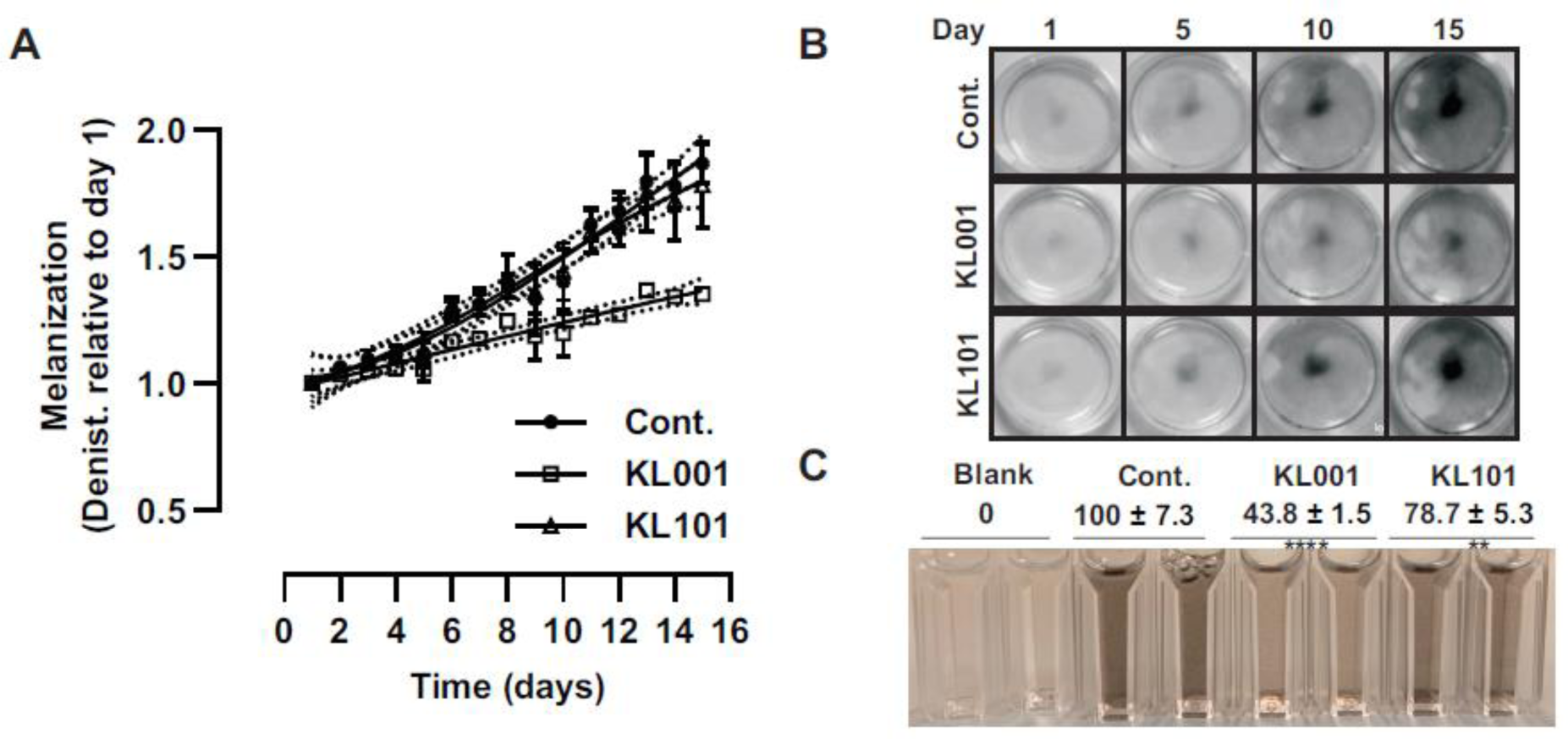
Pharmacological stabilization of Crys inhibit melanin synthesis. **A**) MEX cells depleted of melanin by PTU treatment were seeded in 35 mm dishes and maintained under light:dark cycles (L:D) 12 h light ON 12 h OFF in the presence of KL001 or KL101 (10 uM). The effect on melanisation was determined by densitometric analysis of pictures obtained every day from the cell monolayers. Data from a representative experiment are presented as mean ± SD; n=4; N=2. **B**) Representative picture of dishes taken at the indicated days. **C)** Cells in suspension (∼3 x 10^5^ cells/ml) after 15 days of the indicated treatments. Melanin content is expressed relative to control. Data are the mean ± 95% CI; n=4; N=2. **, p<0.01; ****; p<0.0001 multiple ANOVA followed by Bonferroni.

## 4 Discussion

The control of skin physiology by light occurs by a combination of a direct response to different wavelengths and indirect mechanism via modulation of the synthesis and release of light-regulated hormones, such as melatonin. In this study, we examined the proliferative and melanisation responses of *Xenopus* melanophores exposed to both light and melatonin, and found that:

1. Skin melanophores exhibit a robust endogenous circadian response, with maximum aggregation of melanosomes occurring at night. Yet, we find dark/light exposure had minimal impact on the aggregation/dispersion of melanosomes in cultured MEX cells.
2. Light did disperse melanosomes, but only in a small population (<10%) of cells, even though a vast majority (∼ 70%) show normal dispersion/aggregation melanosome responses to αMSH and melatonin, respectively. The fact that MEX cells respond much more robustly to the neuroendocrine hormones than they do to light agrees with our previous *in vivo* data [8, 40], and suggest that the circadian changes in skin pigmentation are likely driven by the hormone melatonin rather than the direct response of the melanophores to light/dark cycles. Indeed, we showed previously that a switch from light to dark produces no detectable change in skin pigmentation *in vivo* when the pineal complex is absent [8], indicating that melatonin secreted by the pineal complex is responsible for the circadian response. These data also suggest that MEX cells are a useful simplified model to assess the direct response of cells to distinct stimuli.
3. MEX cells respond to physiological melatonin concentrations by reduced proliferation and melanin synthesis, as do rodent [52] and human [12, 18, 44] melanocytes, suggesting the responses are conserved during the evolution. Our interest in analyzing melanophores from amphibians stems from their location on light-exposed integuments, as is the case in fish and reptiles, but in contrast with mammals and birds where they are protected by fur and feathers, respectively. All vertebrates produce and release melatonin during the dark phase, independently of their diurnal or nocturnal behavior. Thus, we predicted that melatonin would elicit similar cellular/physiological responses as darkness. Indeed, melatonin and dark conditions both cause melanosome aggregation and reduce the melanisation of melanophores. Interestingly, however, while melatonin reduces melanophore proliferation, darkness promotes cell division, arguing that melanophore proliferation s and the circadian cycle although regulated by light/dark are mechanistically independent process.
4. Thus, while both proliferation and melanisation are regulated by light, our data suggest that only melanisation is driven by the circadian rhythm; First, exposing cells to continuous dark or continuous light, two well know methods to alter circadian cycle, inhibit melanin synthesis. Second, the expression of all the clock genes analyzed (*per1*, *per2*, *per3*, *cry1*, *cry2*, *cry4*, *clock*, and *bmal1*) is similar for MEX cells in middle of the dark phase vs. those that are melatonin-treated and analysed in the middle of the light phase. While overexpression of individual Crys does not impact melanisation, we found KL001, which prevents the ubiquitin-dependent degradation of both CRY1 and CRY2 [50, 51], strongly reduces MEX cell melanisation. A similar effect of KL001 is seen for B16 melanoma cells [49]. Through its actions on Cry1/2, KL001 lengthens the circadian rhythm of zebrafish [53]. Thus, our data argue that melanisation requires Cry coordination of the circadian clock. Cry1 and Cry2 may need to work together to impact melanisation, in that only the double knockout of both *Cry1* and *Cry2* affects melanisation of murine hair follicles [19]. In agreement, we find KL101, which stabilizes CRY1 but not CRY2 [51], only partially reduces melanisation. The fact that silencing BMAL or PER1 increases melanin pigment in human hair follicles supports the idea that disrupting the circadian cycle alters melanisation [27]. Note that in mammals, CRY1 and CRY2 have redundant functions in clock gene regulation in that degradation of both CRYs simultaneously activates CLOCK-BMAL1, allowing the system to restart a new 24-h cycle [54]. CRY1 and CRY2 can also have distinct actions [54, 55], as we observed with overexpression of *cry2* but not *cry1* altering the mRNA levels of other *cry* genes.
5. We observed that *cry1* and *cry2* mRNA levels in MEX cells are highest during the dark phase, as is the case for the *Xenopus laevis* retina [56] and Rana bullfrog brain [57]. These data suggest that in amphibians *cry* expression is regulated in a coordinated manner within the skin and the two light-sensitive neuronal oscillators (brain/pineal complex and the eye). We also find melatonin promotes *cry1* and *cry2* skin expression. While this mechanism is not yet described for mammalian skin, melatonin induces a strong and transient expression of Cry1 in the pars tuberalis of the pituitary gland, a tissue with the same embryonic origin as skin melanophores [58]. The regulation of *cry1* and *cry2* by melatonin is likely complex in that CRY1 and CRY2 themselves regulate mouse pineal melatonin synthesis via circadian and photic signals from the SCN, a mechanism lost in *Cry1/2* double knockout animals [55]. The fact that applying external light to the skin of these knockout animals fails to restore core gene expression [19] supports the idea that a neuroendocrine (melatonin) mechanism is necessary for the circadian physiology of the skin in rodents, as our data suggests in *Xenopus*.

The current understanding is that Cry1 and Cry2 function as transcriptional regulators with low FAD binding affinity, while Cry4 exhibits high affinity for FAD. This latter property enables Cry4 to serve both as a UV/blue light photoreceptor and a light-dependent magnetosensor, critical for bird migration [29, 31, 32]. Our results show that in contrast to *cry1* and *cry2*, the expression of *cry4* mRNA did not vary in response to either light/dark cycles or melatonin treatment. Similarly, light does not affect *cry4* expression in the zebra finch (Taenopygia guttata) [59] and European robin (Erithacus rubecula) [32] retina. Notably, *cry4* disappeared during mammalian evolution (Synapsids) but remains in Sauropsids (birds and reptiles) as well as in the amphibian lineage, predating the sauropsid/synapsid split [30]. Loss of function studies will be necessary to understand the specific role of Cry4 on proliferation and/or skin pigmentation.

In conclusion, we analyze and compare the impact of the light/dark cycle and the associated neuroendocrine hormone melatonin on the growth and melanisation of *Xenopus* melanophores. We propose that melatonin serves as a more potent inducer of cell aggregation than darkness *in vivo*, and we find that this differential response extends to melanophore behavior *in vitro*, possibly indicating conservation of the response across organisms with exposed and unexposed integuments. Notably, we observe contrasting effects on cell duplication of the dark condition and melatonin, with the former decreasing and the latter increasing population doubling time. Yet, both conditions upregulate the expression of *cry1* and *cry2*. Interestingly, overexpression of these *crys* individually does not alter the proliferation and pigmentation responses of melanophores. Instead, we find external cues (continuous light, dark or melatonin) or pharmacological deregulation of the circadian rhythm alter melanin synthesis. These findings underscore the complexity of melanophore regulation and suggest avenues for future research aimed at unraveling the intricate mechanisms governing melanophore behavior in response to environmental cues.

## Acknowledgements

This work was supported by an operating grant from the Natural Sciences and Engineering Research Council of Canada (NSERC) to SM. We thank Ms. Carrie Hehr for excellent technical assistance.

## Author contributions

GEB, ND and NH designed the experiments. GEB, ND, NH and RB performed the experiments and analyzed the data. HZ-M and CS provided conceptual advice and critically revised the manuscript. GEB and SM supervised the study and wrote the paper.

---

**Supplementary Figure 1.**
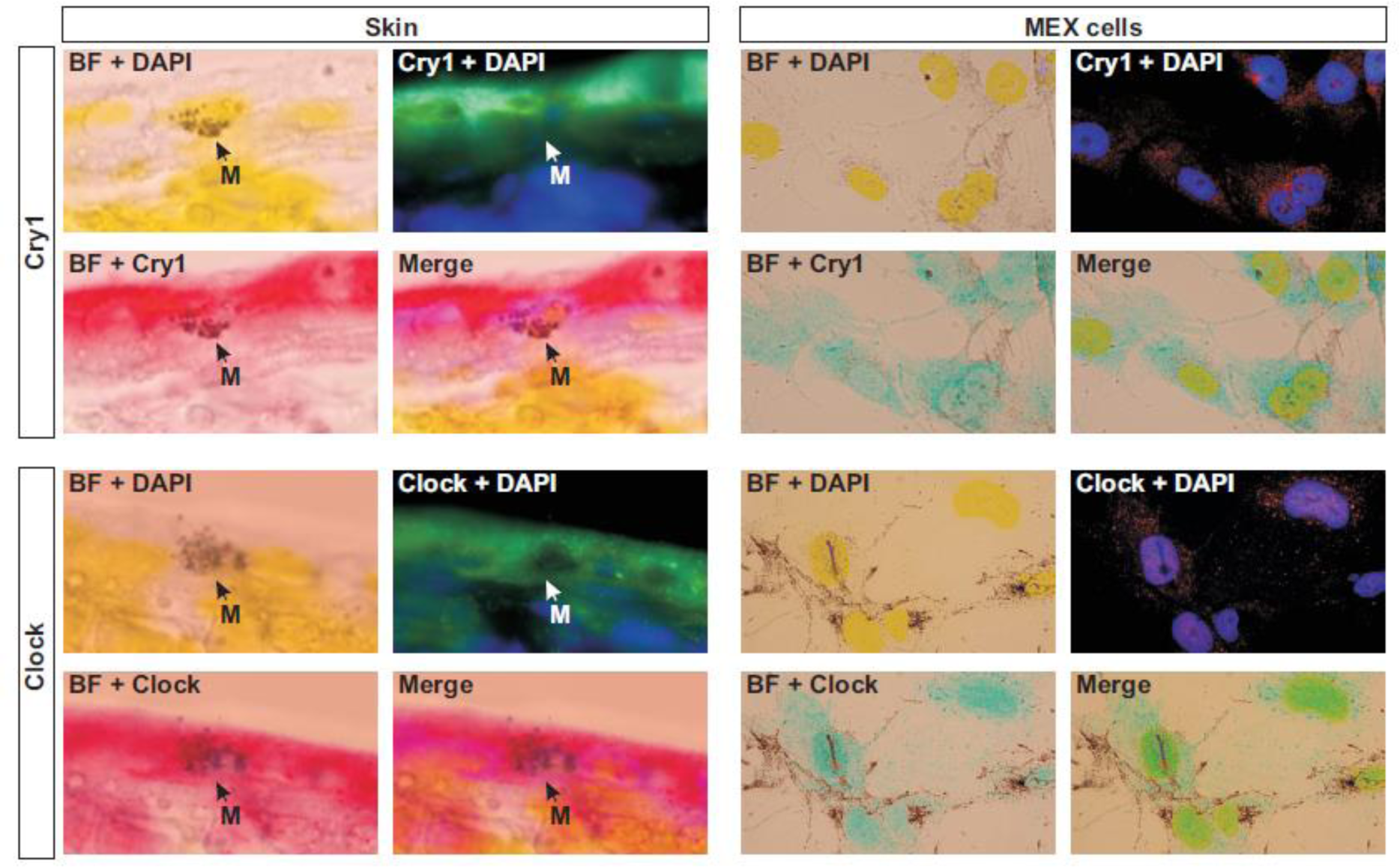
Immunohistochemistry for Cryptochrome 1 (Cry1) and Clock in the skin of sections of stage 43/44 tadpoles (left) and for MEX cell cultures (right). Bright field (BF) shows melanosome granules while DAPI staining (blue) labels cell nuclei. Cry1 and Clock are expressed in melanophores *in vivo* (skin) and *in vitro* (MEX cells).

**Supplementary Figure 2.**
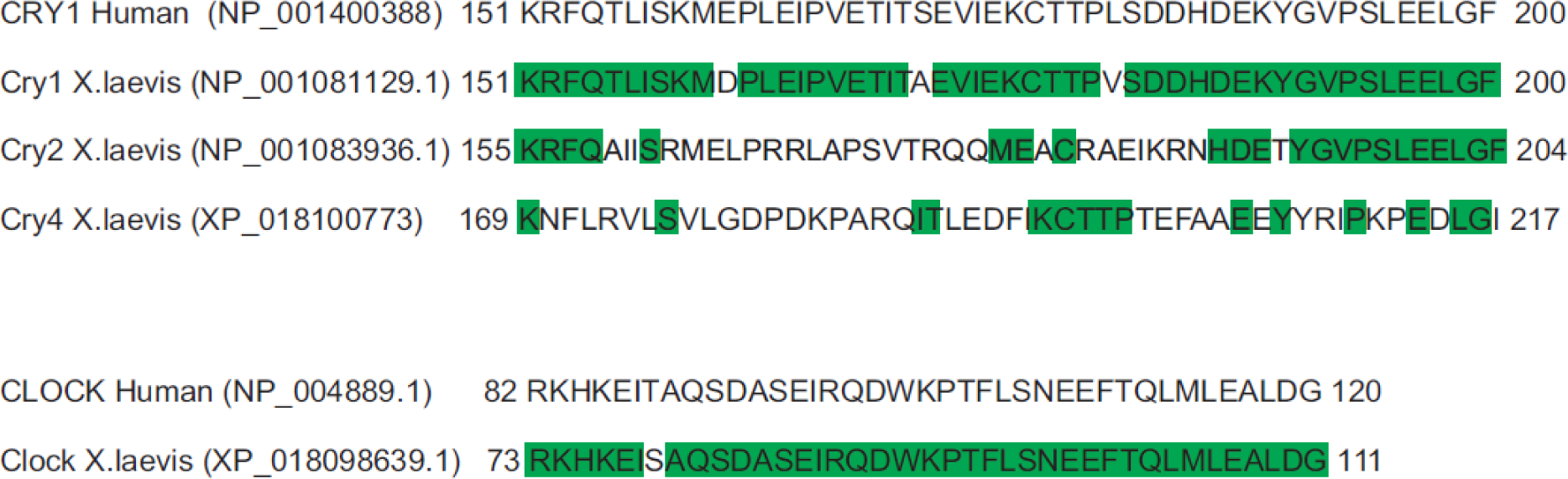
Alignment of the amino acid sequences of human CRY1 (top) and CLOCK (bottom) used to generate antibodies against the *X. laevis* Crys (Cry1, Cry2 and Cry4) and Clock. Highlighted green denotes identical amino acids between species.

